# Cholinergic neuromodulation of prefrontal attractor dynamics controls performance in spatial working memory

**DOI:** 10.1101/2024.01.17.576071

**Authors:** Alexandre Mahrach, David Bestue, Xue-Lian Qi, Christos Constantinidis, Albert Compte

## Abstract

The behavioral and neural effects of the endogenous release of acetylcholine following stimulation of the Nucleus Basalis of Meynert (NB) have been recently examined (Qi et al. 2021). Counterintuitively, NB stimulation enhanced behavioral performance while broadening neural tuning in the prefrontal cortex (PFC). The mechanism by which a weaker mnemonic neural code could lead to better performance remains unclear. Here, we show that increased neural excitability in a simple continuous bump attractor model can induce broader neural tuning and decrease bump diffusion, provided neural rates are saturated. Increased memory precision in the model overrides memory accuracy, improving overall task performance. Moreover, we show that bump attractor dynamics can account for the nonuniform impact of neuromodulation on distractibility, depending on distractor distance from the target. Finally, we delve into the conditions under which bump attractor tuning and diffusion balance in biologically plausible heterogeneous network models. In these discrete bump attractor networks, we show that reducing spatial correlations or enhancing excitatory transmission can improve memory precision. Altogether, we provide a mechanistic understanding of how cholinergic neuromodulation controls spatial working memory through perturbed attractor dynamics in PFC.

**Significance statement:** Acetylcholine has been thought to improve cognitive performance by sharpening neuronal tuning in prefrontal cortex. Recent work has shown that electrical stimulation of the cholinergic forebrain in awake-behaving monkeys induces a reduction in prefrontal neural tuning under stimulation conditions that improve performance. To reconcile these divergent observations, we provide network simulations showing that these derive consistently from specific conditions in prefrontal attractor dynamics: firing rate saturation leads to increased storage precision and reduced neural tuning upon cholinergic activation via an increase in neural excitability, a reduction in neural correlations, and an increase in excitatory transmission. Our study integrates previously reported data into a consistent mechanistic view of how acetylcholine controls spatial working memory via attractor network dynamics in prefrontal cortex.

## Introduction

Understanding how neuromodulatory systems affect associative cortex dynamics during cognitive function is fundamental to acquiring insights into cognitive control processes. Such understanding is also essential for advancing hypotheses for the network mechanisms underlying neuropsychiatric disorders, often characterized by subtle cognitive processing alterations. In this regard, working memory (WM) is of particular relevance. For one, WM is known to be modulated by the dopaminergic, noradrenergic, and cholinergic systems (Cools and Arnsten, 2021). Besides, it is closely associated with network activity in the prefrontal cortex (PFC), for which computational models have successfully linked network dynamics with behavior (Compte et al., 2000; Wimmer et al., 2014; Inagaki et al., 2019; Barbosa et al., 2020). Furthermore, WM is affected in most neuropsychiatric disorders (Forbes et al., 2009). However, the causal chain from cellular neuromodulation to changes in network dynamics and behavioral performance in WM remains largely unknown, even for simple behavioral readouts, such as spatial memory precision in delayed-response tasks (Stein et al., 2021).

A series of recent studies have investigated how the release of endogenous acetylcholine through the stimulation of the nucleus basalis (NB) of Meynert impacts cognitive functions, including spatial and color WM and sustained attention (Blake et al., 2017; Liu et al., 2017; Qi et al., 2021). Remarkably, they reported that NB stimulation leads to better behavioral performance, an uncommon observation upon interference with neuromodulatory systems. In addition, they investigated the neural correlates of improved performance in PFC (Qi et al., 2021). Unexpectedly, they found that NB stimulation induced a general increase in excitability and a widening of neuronal tuning curves and memory fields. A decrease in selectivity has been observed due to cholinergic stimulation applied iontophoretically with high doses of cholinergic agonists (Major et al., 2018; Vijayraghavan et al., 2018; Galvin et al., 2020b). However, this effect has been assumed to correspond to the descending section of an inverted-U function, representing a regime over which cholinergic agonists impair performance (Galvin et al., 2020a). The effectiveness of drugs targeting other neurotransmitter systems, e.g., dopamine, is often interpreted as increased stimulus selectivity (Williams and Goldman-Rakic, 1995). Thus, dopamine agonists are known to “sculpt” neuronal activity and improve spatial selectivity at low doses, which is assumed to confer the beneficial effect of dopamine in behavior (Vijayraghavan et al., 2007) as the result of more efficient coding (Fitzpatrick et al., 1997). The concurrent improvement in performance and broadening of tuning raises the question of how weaker memory code at the neural level can lead to better behavioral performances. An answer to this question requires a mechanistic understanding to link PFC dynamics with behavior consistently.

Bump attractor models (Wilson and Cowan, 1972; Amari, 1977; Compte et al., 2000) have successfully explained complex aspects of behavior in spatial WM tasks (Wimmer et al., 2014; Barbosa et al., 2020). In such models, a neural population represents a continuous feature by localized elevated activity (bump attractor) self-sustained through the mnemonic delay period. Continuous bump attractor dynamics are usually investigated in idealized settings with network homogeneity and perfectly symmetric connectivity. However, their more plausible biological implementations must deal with inhomogeneities and asymmetries that can significantly alter the dynamical features of the bump attractor (Renart et al., 2003; Hansel and Mato, 2013; Seeholzer et al., 2019; Darshan and Rivkind, 2022).

Here, we first use a homogeneous, symmetric firing-rate bump attractor network model (Amari, 1977) — which we will refer to as the continuous bump attractor model— to build a mechanistic understanding of how increased excitability in the network can explain the effects of NB stimulation in PFC and its behavioral outcomes in monkeys engaged in a spatial WM task (Qi et al., 2021). We show that bump attractor models can combine broader neural tuning during the delay and improve overall performance in the task, provided that neurons undergo firing rate saturation during the dynamics. Behavioral improvement is due to a reduction in bump diffusion, confirmed in the experiments’ behavioral data.

Finally, we investigate how these ideal attractor-like qualities can emerge in more realistic, heterogeneous, and asymmetric networks of spiking neurons (discrete bump attractor networks) (Hansel and Mato, 2013). We find that heterogeneity per se does not alter our conclusions. Still, when considering network dynamics of strongly recurrent excitatory and inhibitory populations, the cholinergic activation in such discrete bump attractor networks must include other mechanisms than just enhanced excitability to reproduce the experimental data, such as reduced network spatial correlations (Thiele et al., 2012; Chen et al., 2015; Minces et al., 2017).

## Materials and Methods

### Electrophysiology

The experimental data included here was previously reported (Qi et al., 2021). Two adult male rhesus monkeys (*Macaca mulatta*) were implanted with a stimulating electrode targeting the anterior portion of the NB (Fig. 1A). Intermittent stimulation of the NB was applied for 15 seconds at 80 pulses per second, followed by approximately 45 seconds with no stimulation (Fig. 1C). The stimulation was applied during the inter-trial interval of the behavioral task (below). In addition, neural recordings were obtained from arrays of 1-4 electrodes over areas 8a and 46 of the dorsolateral PFC and were previously reported by Qi et al. (2021).

**Figure 1.**
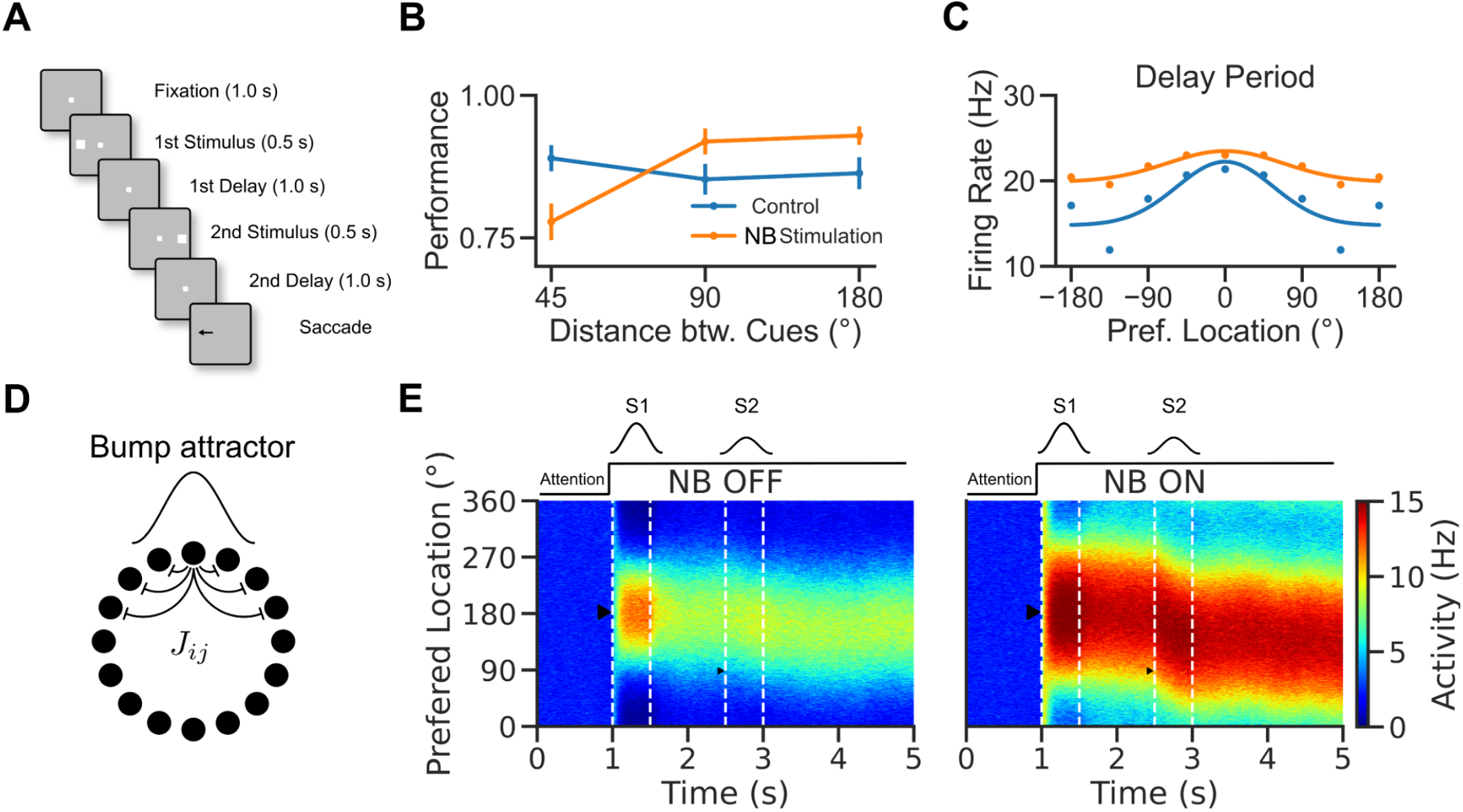
A rate model to reconcile cholinergic modulation of behavior and neural tuning in an ODR task. A. Visuospatial WM task with distraction (Qi et al., 2021). B. Performance of the monkeys in control and NB-stimulated trials. Performance was measured as percent of trials with responses within 7º visual angle of the target stimulus. C. Prefrontal neuron location selectivity in the delay period (adapted from Qi et al., 2021). Blue: control trials. Orange: NB-Stimulated trials. Dots: data. Lines: fits. D. Network scheme. The network consists of a population of 1000 rate units with tuned connections. E. Activity vs. neurons vs. time, left: NB OFF (*I*_0_ ^*OFF*^ = 10*Hz*) condition, right: NB ON condition (*I* _*O*_ ^*ON*^ = *I* _*O*_ ^*OFF*^ + 20*Hz*, simulates increased excitability induced by NB stimulation). After 1s in baseline activity, neurons receive a tuned stimulus, S1, and a nonspecific attention signal is switched on. The activity becomes structured into a bump. The bump is only slightly perturbed by a second weak stimulus, S2, so the bump maintains a location around S1. Triangles show the locations of the stimuli.

### Task

Monkeys were trained to perform a variation of the oculomotor delayed response task, where two visual stimuli appeared in sequence (Fig. 1B). They learned to remember the location of either the first (Remember 1st) or the second (Remember 2nd) stimulus and make an eye movement to its location depending on the color of the fixation point (white remember 1st/blue remember 2nd). The task was distributed in blocks. Remember 1st, blocks include retrospective distractors (monkeys had to remember the first and ignore the second). Remember 2nd, blocks contained prospective distractors (monkeys had to ignore the first and remember the second stimulus). Trial blocks included two null conditions in which either the first or the second stimulus is omitted (Remember 1st/absent 2nd and Remember 2nd/absent 1st).

After a 1s fixation period, the first stimulus was presented for 0.5s. After a 1s delay (delay 1), the second stimulus was presented for 0.5s. Monkeys had to respond with a saccade after another 1s delay (delay 2). The monkeys were rewarded with juice after reporting the correct location. Gaze deviation beyond the fixation window led to immediate trial termination without reward.

Each stimulus was displayed at one of eight locations arranged along a circular ring (separated by 45 degrees). The angular distance between the target stimulus and the distractor could be 0, 45, 90, or 180 degrees. Additional details regarding the task structure can be found in the original article (Qi et al., 2021).

### Continuous bump attractor network

Our first model consists of a homogeneous bump attractor network of a single population of rate neurons, similar to that proposed by (Amari, 1977).

#### Rate dynamics

The equations that define the evolution of the rates of the neurons in our model are:

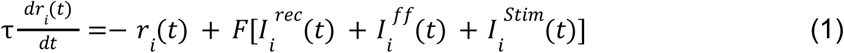

where we fixed, τ = 20*ms*.

Except for Figure 2A-C where we consider, a threshold linear transfer function

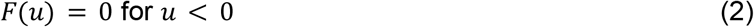

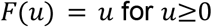

we will assume that the input/output neural transfer function is a sigmoid-like function that saturates for large input values:

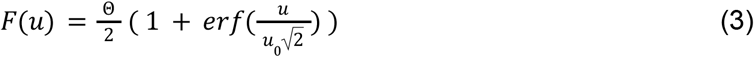

where, Θ= 15*Hz*, is the saturation threshold of the neural activity, and μ_0_ = 1*Hz*.

**Figure 2.**
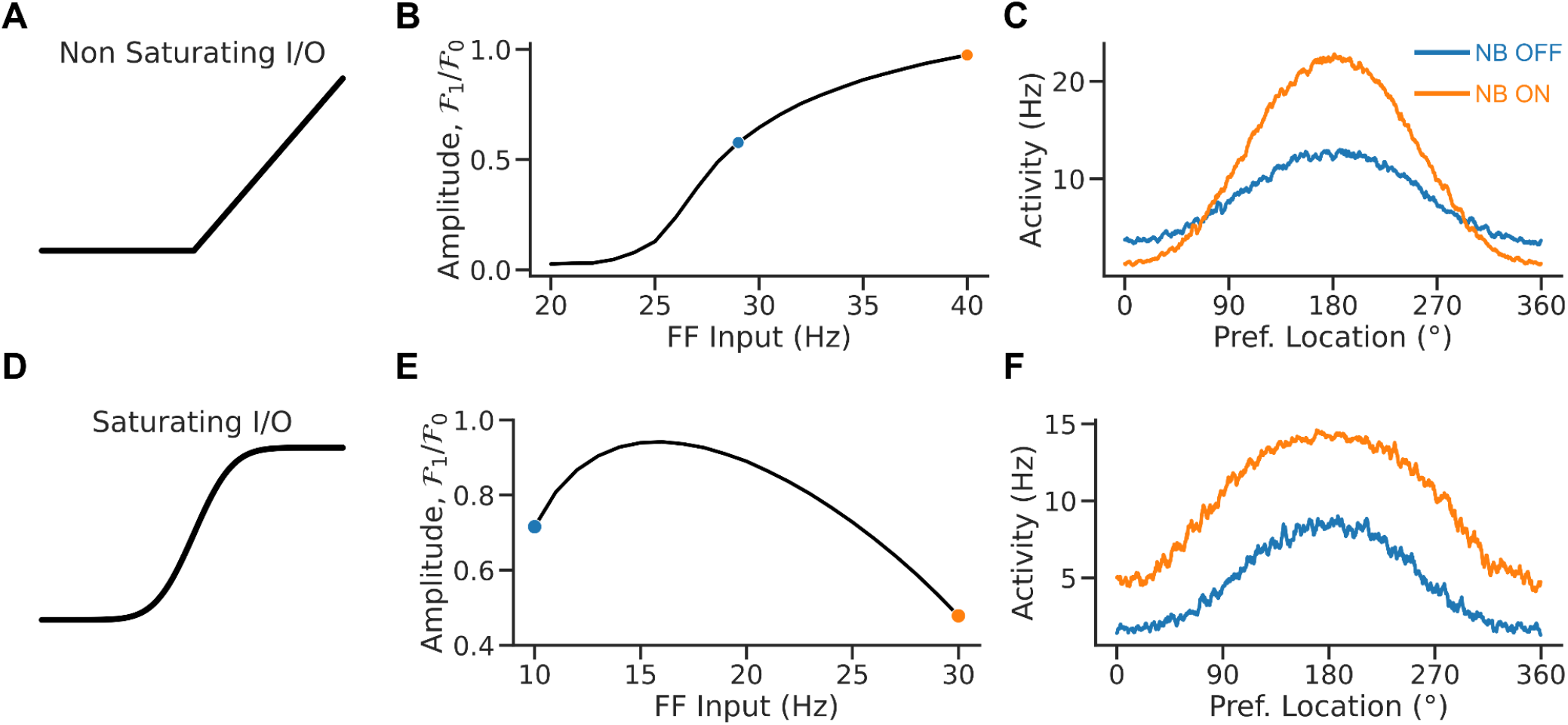
Rate saturation accounts for the broadening tuning following NB stimulation. A. Non-saturating I/O function. B. Relative bump amplitude (i.e., the ratio of Fourier modes F1/F0 of the population rate vector, see methods) vs. the strength of the feedforward (FF) input. Tuning sharpens with stronger FF inputs. C. Population tuning curve for NB ON and NB OFF conditions (matching colors in B). D. Saturating I/O function (saturation occurs at 15*Hz*). E. Relative bump amplitude vs. FF input. Tuning broadens with stronger FF input values. F. Population tuning curve for NB ON and NB OFF conditions (matching colors in E).

#### Connectivity

The network consists of *N* = 10_00_ fully interconnected neurons. Neurons lie on a ring and present selectivity to stimulus locations (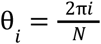 for *i*∈[1, *N*]). Neurons with similar preferred locations are more strongly connected than neurons coding for distant locations.

We model this with a connectivity profile that follows a cosine shape,

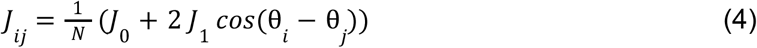

In the simulations, we chose: *J*_0_ =− 2. 75 and *J*_1_ = 1. 1, therefore all connections are negative.

In the case of the heterogeneous bump attractor network, we add an extra noisy term to the connections,

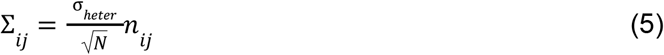

where *n* _*ij*_ are random and independently distributed values with zero mean and unit variance.

#### Recurrent inputs

The recurrent input to neuron *i* is given by,

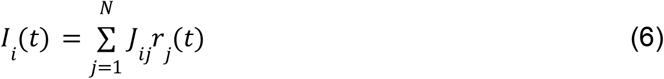

#### Feedforward input

Neurons in the network receive an external input that is a feedforward input with mean, *I*_0_, and temporal variance, η_0_,

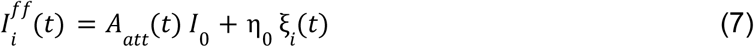

where we introduce an attention switch, *A* _*att*_ (*t*), which turns on (*A* _*att*_ (*t* > *t* _*S*1_) = 1 and *A* _*att*_ (*t* ≤ *t* _*S*1_) = 0) upon stimulus presentation that remains on for the rest of the simulation. In the simulations, we took: *I*_0_ = 10*Hz* and η_0_ = 5. 48*Hz*.

#### Stimulus input

During stimulus presentation, neurons receive an additional tuned input,

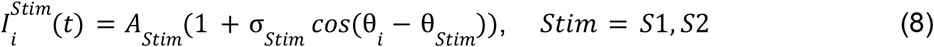

where *A*_*Stim*_ is the strength of the stimulus and σ_*Stim*_ is the stimulus modulation and θ_*Stim*_ the stimulus location. In the simulations, we took: *A* _*S*1_ = 1. 0*Hz*, σ_*S*1_ = σ_*S*2_ = 1. 0.

Moreover, to simulate the effect of distraction in Remember 1st trials, we assume that *A*_*S*2_ is drawn from a truncated Gaussian distribution with mean.0 05*Hz* and standard deviation.0 75*Hz*, the negative values of *A*_*S*2_ are set to zero.

#### Tasks

To reflect differences in top-down influences in the specific task blocks, we used different stimulus parameters to simulate the *Remember 1st* and *Remember 2nd* experimental task conditions. Specifically, the second stimulus was weaker in *Remember 1st* than in *Remember 2nd*.

#### NB stimulation

Since the release of acetylcholine in PFC is thought to result in the depolarization of excitatory pyramidal neurons through the blockade of hyperpolarizing currents (McCormick and Prince, 1986; Delmas and Brown, 2005; Gulledge and Stuart, 2005; Carr and Surmeier, 2007; Zhang and Séguéla, 2010), we modeled NB stimulation as an increase of the constant external input to the excitatory neurons, *I*_0_ ^*ON*^ = *I*_0_ + δ*I*_0_. In the simulations, we selected: δ*I*_0_ = 20*Hz*. With this parametric choice, baseline firing rate went from 4.8 Hz to 10 Hz with NB stimulation (compared with 10.9 Hz to 13.4 Hz in the experimental data of Qi et al. 2021).

#### Bump Dynamics

In analyzing the simulation results of the different models, we measure changes in the network selectivity by referring to the relative amplitude of the bump. We define it as the ratio between the first and zero moments of the discrete Fourier transform of the rate population vector:

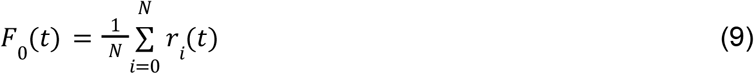

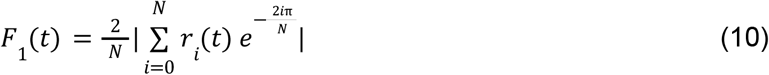

where |.| is the modulus operator.

Moreover, we compute the center of mass of the bump as:

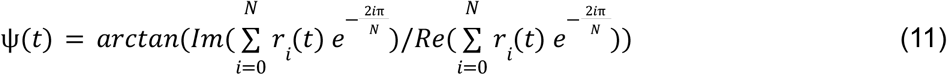

and we compute population tuning curves by smoothing the population rate vector, *r*_*i*_(*t*) with a rolling average over nearby neurons.

#### Model performance

In the attractor rate model, errors in the model can have two components:

1. Distraction and heterogeneities systematically bias the bump center endpoint affecting the accuracy of the response. By computing the bump center endpoint deviation from S1 averaged over trials, we can estimate this bias as

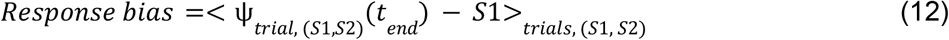 In the continuous attractor model with no distraction, there are no response biases. However, in trials with distraction (Fig 5), distraction shifts the response in a distance-dependent manner. We define a measure for this shift in a signed manner as

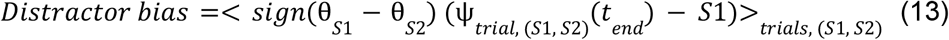
2. The diffusion of the bump on the ring around the location of the stimulus affects the precision of the response. By computing the bump center endpoint deviation from its mean over trials,

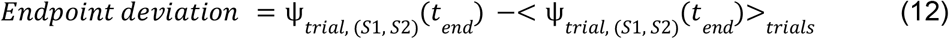

we can define the average variance in model responses by

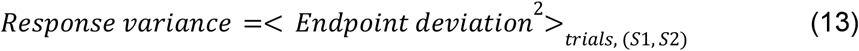 The average standard error in responses as

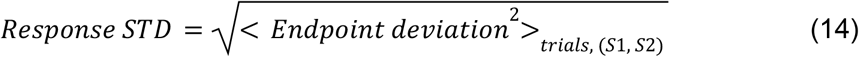

and define the diffusivity of the random process as

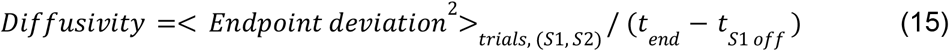 Note: in figure 4, we estimate response bias and STD for model responses within 30° of S1.

**Figure 3.**
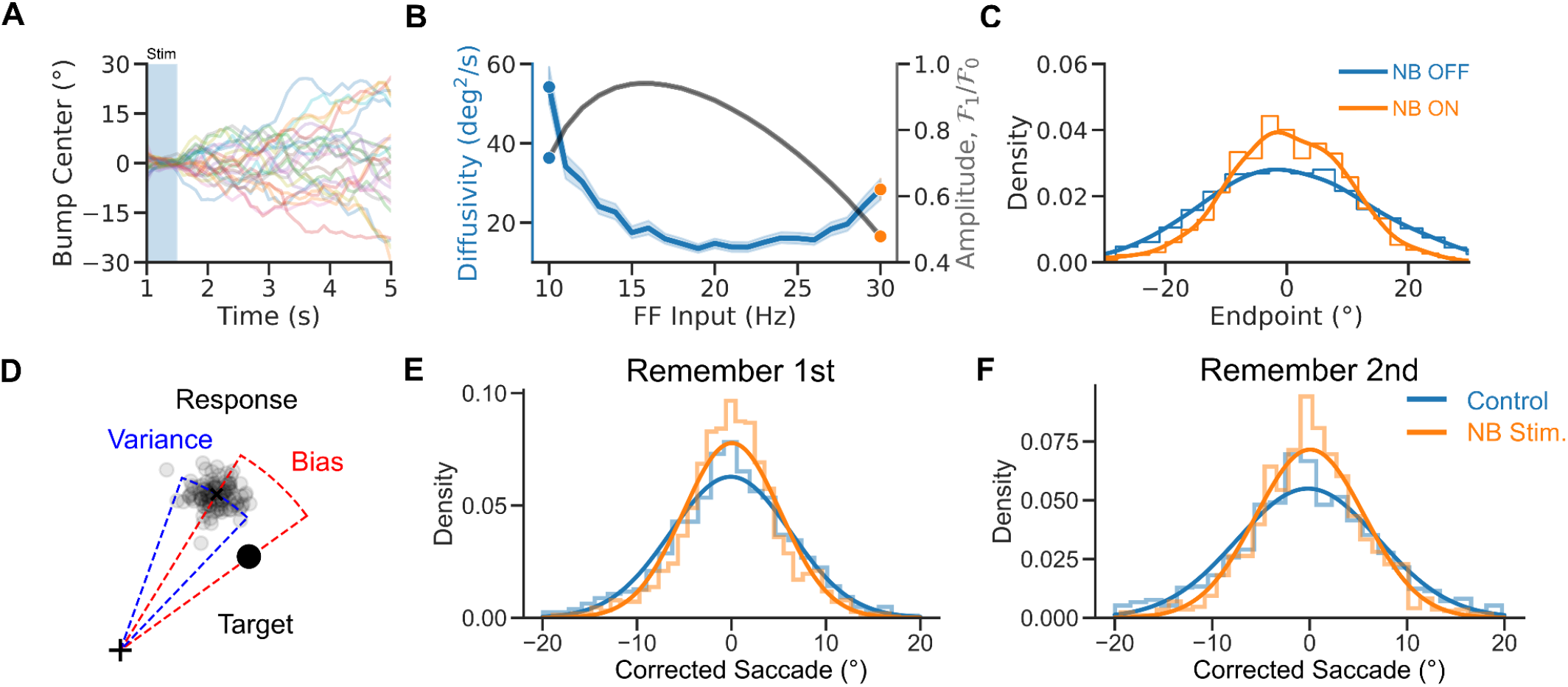
A diffusion process underlies NB stimulation-induced improved performance. A. Bump center vs. time for 50 network initializations (for a cue presented at 0 degrees). The bump center follows a random walk. B. Diffusivity vs. FF input. Diffusion decreases for strong enough FF inputs. Shaded area: bootstrapped 95% confidence intervals. Grey line, relative bump amplitude (see Materials and Methods). C. Distributions of bump centers endpoint location for two different values of the FF input (for 1000 initial conditions). Lines are kernel density estimates. D. Two sources of errors in the experimental data: response bias and response variance. E-F. Distributions of saccades (i.e., mean corrected saccadic endpoint, see Materials and Methods) in Remember 1st and Remember 2nd trials in the data. NB stimulation leads to a sharper distribution of endpoints (see regression analysis, Materials and Methods). Line are Gaussian fits. Blue: control. Orange: NB stimulated trial.

**Figure 4.**
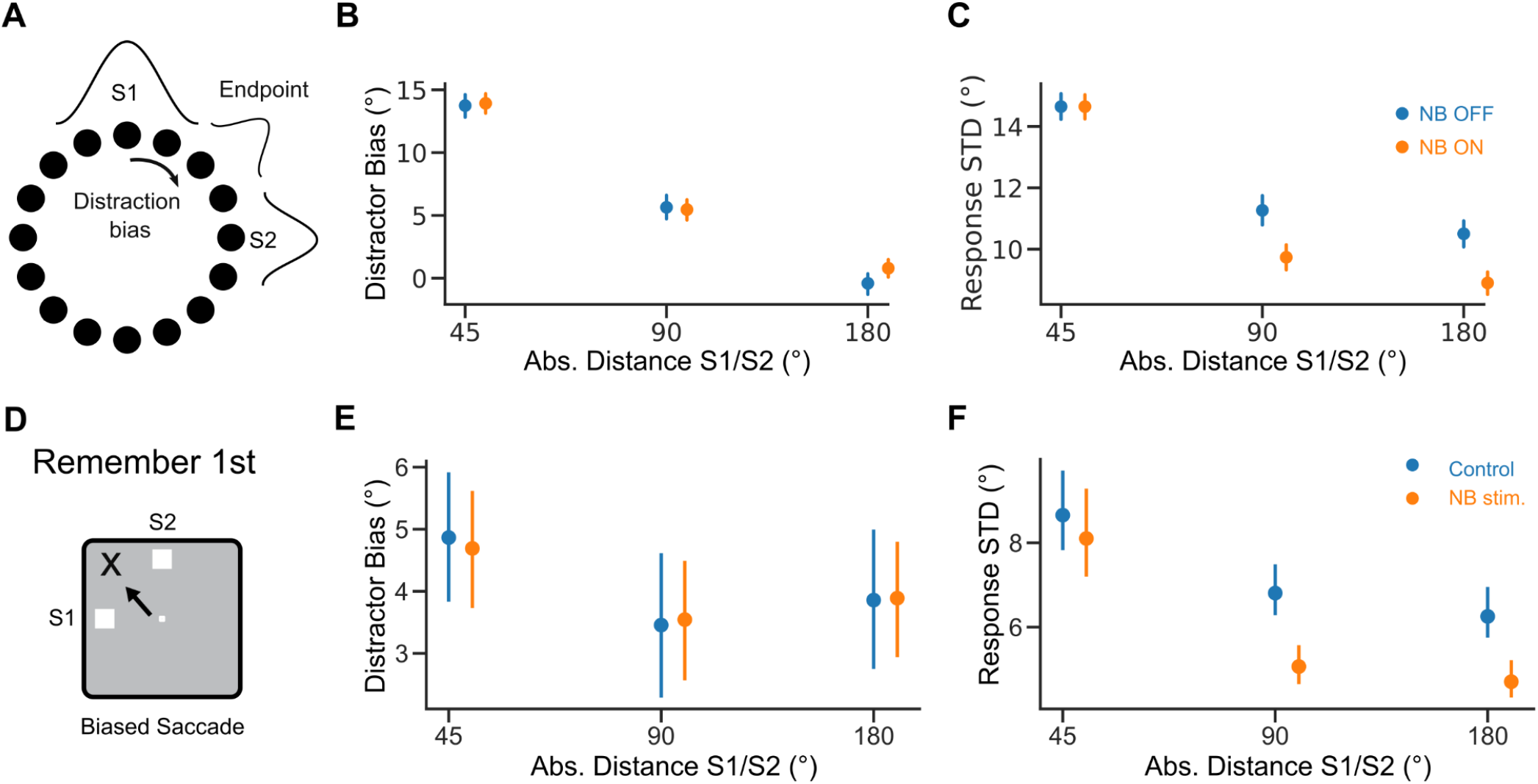
Distraction effect on response variance is distance dependent. A. Distraction by S2 shifts model responses (Distraction Bias). B. Distraction bias vs. absolute distance between S1 and S2. C. Response standard deviation (STD), computed as the square root of the response variance, vs. absolute distance between S1 and S2. Blue: NB OFF condition (*I*_0_ = 10*Hz*). Orange: NB ON condition (*I*_0_ = 30*Hz*). D. Distractor shifts responses in the task. E. Distraction bias vs. absolute distance between S1 and S2 in Remember 1st trials in the data. F. Same but for response STD. Error bars 95% bootstrapped confidence interval. Orange dots are shifted on the x-axis for visibility.

**Figure 5.**
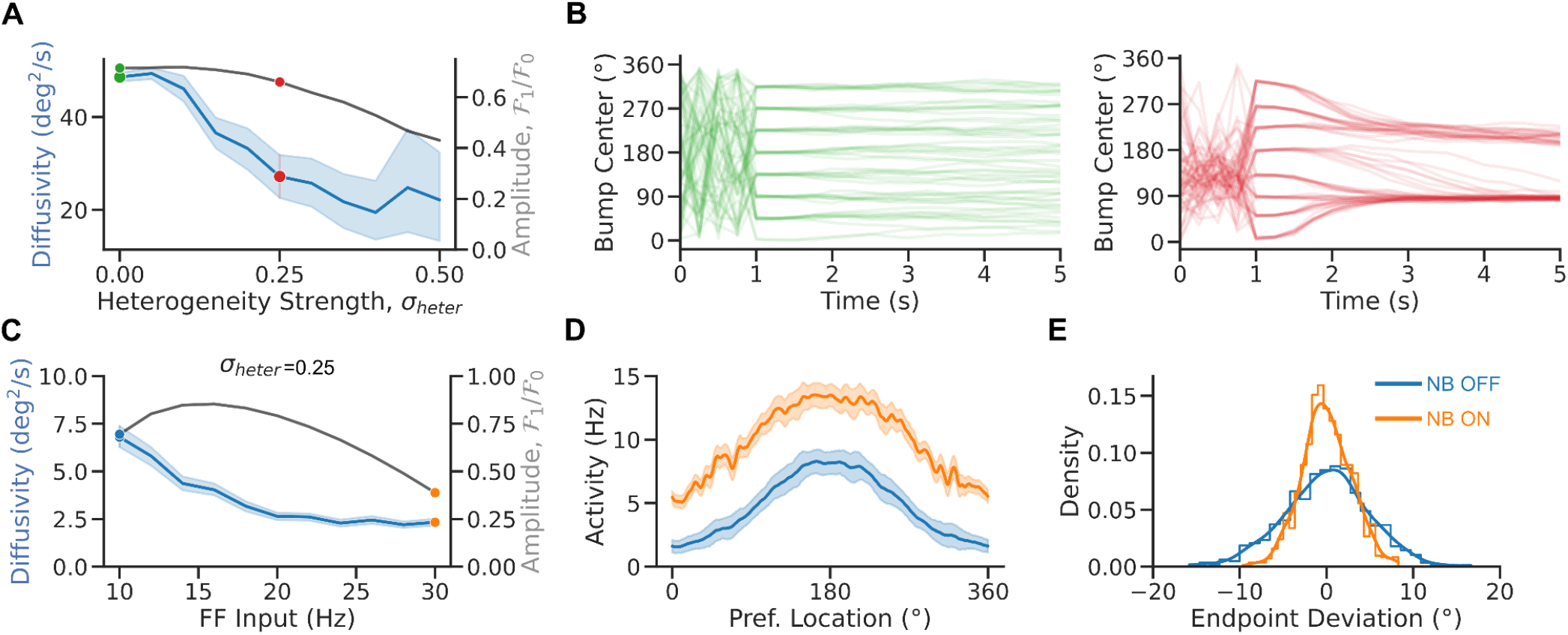
Heterogeneities impair bump diffusion. A. Diffusivity vs. heterogeneity strength (averaged over 8 cues and 25 initial conditions per cue and 100 network realizations). B. Bump center vs. time in the homogeneous (left panel) and heterogeneous (right panel) bump attractor network (for 8 cues and 10 initial conditions per cue for one network realization). C. Diffusivity (averaged over 8 cues and 250 initial conditions per cue, blue) and bump amplitude (black) vs. FF input in the heterogeneous network model (averaged over 8 cues and 125 initial conditions per cue and one network realization, σ _*heter*_ = 0. 25, see Materials and Methods). D. Population tuning curve for NB ON and NB OFF (matching colors in C). E. Bump endpoint deviation (i.e., mean corrected endpoint) for NB ON and NB OFF. Lines: kernel density estimates. Error bands 95% confidence intervals.

### Quantitative and statistical analysis of saccadic errors

#### Response bias and variance

In the data, we decompose saccades in a similar manner. For each monkey, we compute a saccade response bias, Δθ _*i*_, as the deviation of the saccade location, θ _*i*_, to the target stimulus averaged over trials with the same stimuli pair,

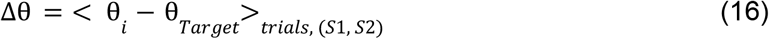

We estimate the saccades response variance by computing the deviation of the saccade to the mean saccade averaged over trials with the same stimuli pair,

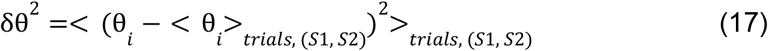

Saccade bias and variance were estimated for saccades within 30° of the reported stimulus.

In figure 3E-F, we tested how task conditions affected saccades response variance in a linear model:

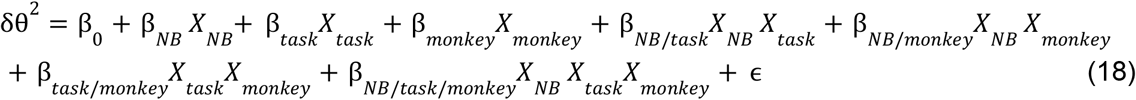

where *X*_*NB*_ = 0 *or* 1 for control or NB stimulated trial, *X* _*task*_ = 0 *or* 1 for Remember 1st or Remember 2nd trial, *X* _*monkey*_ = 0 *or* 1 for each monkey, and ϵ is assumed to be a Gaussian random variable.

We fitted the model’s coefficients with an Ordinary Least Square scheme. We did not find significant interactions between NB and the monkey factor, so we only included the main factor, “monkey”, in subsequent analyses. In the main text, we report the coefficients and p-values of the interaction between NB stimulation and task (β _*NB/task*_).

#### Distraction bias

In our analysis of the effect of distraction on response accuracy (Fig 4), we tested the dependency of distractor biases with the absolute distance between S1 and S2 with the following linear model:

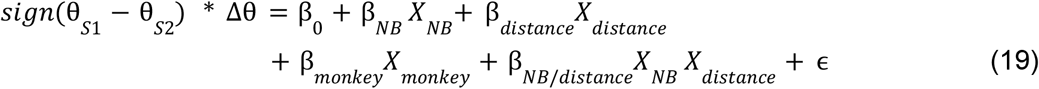

where *X* _*distance*_ = 0 *for* |θ _*S*1_ − θ _*S*2_ | = 45 *deg* and *X* _*distance*_ = 1 *for* |θ _*S*1_ − θ _*S*2_ | = 90 *or* 18_0_ *deg*.

Moreover, we tested the dependency of response variance with the distance between stimuli with the following linear model:

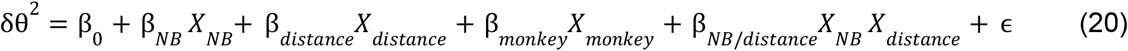

We fitted the coefficients of the model with an Ordinary Least Square scheme. We report in the main text the coefficient and p-value of the interaction between NB stimulation and distance (β _*NB/distance*_).

### Discrete attractor spiking network

Our more realistic spiking model consists of a two-population network of strongly recurrent leaky integrate-and-fire neurons.

#### Leaky integrate-and-fire neuronal dynamics

The dynamics of the membrane potential of the i-th neuron in population A, (*i, A*), is given by

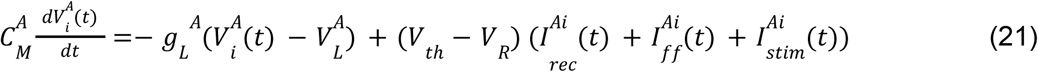

where 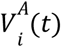 is the membrane potential, 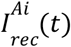 the recurrent input, 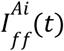 the feedforward input and 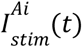 the stimulus input into neuron (*i, A*).

Reset condition: if at time *t* the membrane potential, 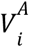, of neuron (*i, A*) crosses the threshold,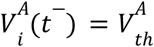, the neuron fires a spike, and its voltage is reset to its resting potential,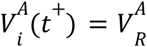.

In the simulations, we chose: 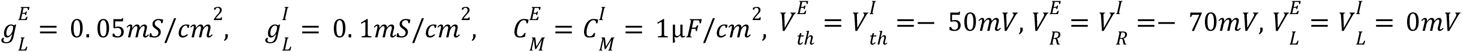

#### Connectivity

The connectivity between population *A* and *B* is a random matrix 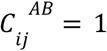 with probability 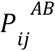 and 0 otherwise. On average, neurons in each population receive inputs from *K*_*B*_ neurons in the presynaptic population *B*. 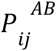 varies with the difference in preferred location between the neurons (Van Vreeswijk and Sompolinsky, 2005),

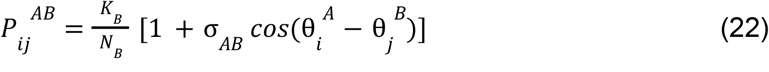

where σ _*AB*_ determines the spread of the projections from population *B* to population *A*. Here 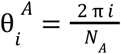_, is_ the preferred location of neuron (*i, A*).

In the simulations, we selected: *N* _*E*_ = 320_00_, *N* _*I*_ = 80_00_, for the number of neurons, *K* _*E*_ = 320_0_, *K* _*I*_ = 80_0_ for the average number of inputs and for the spread of the projections σ _*EE*_ = σ_*IE*_ = σ_*II*_ = 1. 0, and we assume I to E slightly less tuned: σ _*EI*_ = 0. 9 (Kerlin et al., 2010; Hofer et al., 2011)

#### Recurrent inputs

The recurrent input to neuron (*i, A*) from its presynaptic neurons in population B is given by

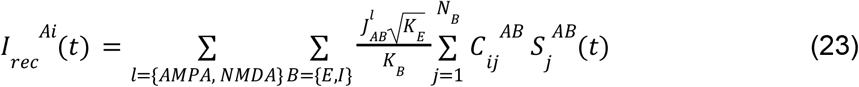

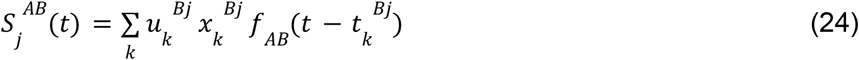

where *t* _*k*_ ^*Bj*^ is the time at which neuron (*j, B*) has emitted its *k* ^*th*^ spike, the sum is over all the spikes emitted by neuron (*j, B*) before time *t*.

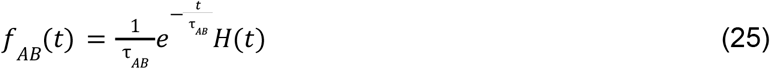

where τ _*AB*_ is the synaptic time constant of the interactions between neurons in population *B* and *A, H* is the Heaviside step function.

In the model, excitatory neurons form a mixture of fast (AMPA) and slow (NMDA) synapses on other excitatory cells and inhibitory neurons.

In the simulations, we took: 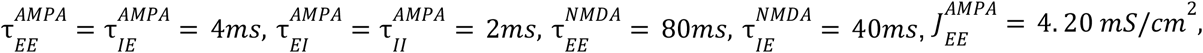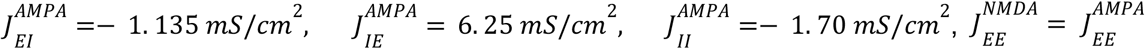 and 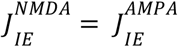

#### Synaptic plasticity

*x*_*k*_^*Bj*^ is the amount of synaptic resources available at the synaptic terminals of neuron (*j, B*) before the spike *t*_*k*_^*Bj*^, and μ _*k*_ ^*Bj*^ is the fraction of these resources used by this spike. The dynamics of these two variables are responsible for the short-term plasticity (STP) experienced by the synapses. We model them as in (Hansel and Mato, 2013):

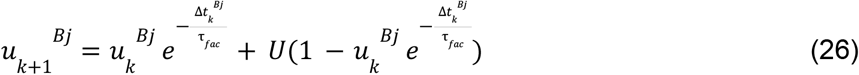

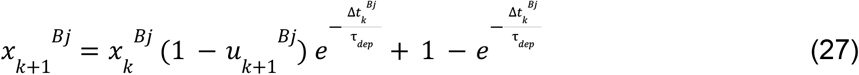

where 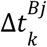 is the interspike interval for neuron (*j, B*) between spike *t* _*k*_ and *t* _*k*−1_. In the simulations, we took: τ _*dep*_ = 25_0_*ms* and τ _*fac*_ = 60_0_*ms*.

#### Feedforward input

At each time point, we assume that the feedforward input, *I*_*ff*_ ^*Ai*^ (*t*), into neuron (*i, A*), is weakly tuned with a random phase *U*(*t*). This leads to the emergence of spatial correlations in the recurrent layer. We model this as

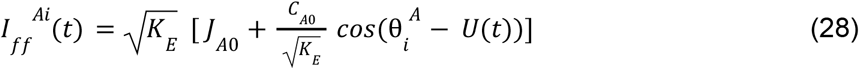

where *J*_*A*0_ is the strength of the external input, at each time step, *U*(*t*) is drawn uniformly between _0_ and 2π, and *C*_*A*0_ controls the strength of the feedforward correlations. In the simulations we choose: *J*_*E*0_ = 1. 95 *mS/cm*^2^, *J* _*I*0_ = 1. 675 *mS/cm*^2^, *C* _*E*0_ = 0. 016, *C* _*I*0_ = 0. 0134 when not mentioned otherwise.

Since correlations in the recurrent input are quite small in strongly recurrent networks (Darshan et al., 2018), most of the correlations in the net input to a neuron come from the feedforward contribution:

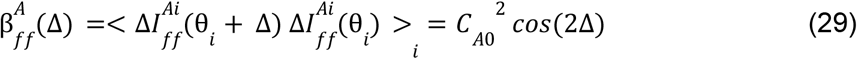

where Δ ∈ [0, 2 π], is a location on the ring.

#### Stimulus Input

During stimulus presentation, the excitatory population receives an additional tuned input,

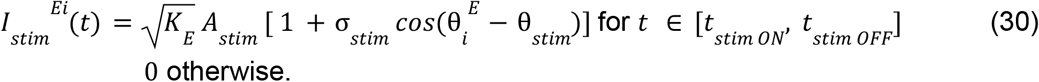

where θ_*stim*_ is the location of the stimulus, *A*_*stim*_ is the strength of the stimulus and σ_*stim*_ is the footprint of the stimulus. The inhibitory population receives no stimulus.

In the simulation, we took *A* _*stim*_ = 0. 1 *mS/cm* and^2^ σ_*stim*_ = 1.

#### Bump Dynamics and model responses

We compute bump amplitude, center, diffusitivity and endpoint deviations in the same way we did for the rate model.

#### NB stimulation

We model NB stimulation’s main effect as an increase of the mean feedforward input to the excitatory neurons, *J* _*E*0_^*ON*^ = *J* _*E*0_ + δ*J* _*E*0_. In Figure 7, we also consider the cases where, in addition to an increase in feedforward input: 1) excitatory synaptic strength, *J* _*AE*_, ∀ *A* ∈ *E, I* increase; 2) inhibitory presynaptic strength, *J* _*AI*_, ∀ *A* ∈ *E, I* decrease; 3) the modulation of the spatial feedforward correlations to the E neurons, *C* _*E*0_ decreases.

## Results

### Homogenous, continuous attractor models

We used numerical simulations to investigate the network mechanisms underlying the reported cholinergic neuromodulation of prefrontal activity in a visuospatial WM task (Qi et al., 2021). In this task (Fig 1A), monkeys were trained to saccade to the location of one of two sequentially presented visual cues following a delay period. In alternating blocks of trials, the color of the fixation dot signaled to the monkey whether a reward would be given for an accurate saccade to the remembered location of the first target S1 (Remember 1st trial) or the second target S2 (Remember 2nd trial). Here, we used a computational approach to reconcile evidence collected during the distractor condition (Remember 1st, Qi et al., 2021). Monkeys showed a general increase in performance in this task following NB stimulation, except for trials when S2 appeared close to S1, where NB stimulation impaired performance (Fig 1B). This general increase in performance by NB stimulation was mirrored at the neural level by erosion of PFC neural tuning to the stimulus (Fig 1C). We developed a simple continuous bump attractor rate model to understand the conditions for this modulation of neuronal tuning by NB stimulation (Fig 1D) (Amari, 1977; Ben-Yishai et al., 1997; Itskov et al., 2011). In the model, following the presentation of a location-tuned stimulus (S1, S2), the network maintains information about the stimulus’ spatial location through a localized and persistent increase in activity (Fig 1E) (Wilson and Cowan, 1972; Compte et al., 2000). We model distraction with a second stimulus weaker than the first on average but varies strongly from trial to trial (see Material and Methods), which simulates variable attentional filtering mechanisms occurring in upstream areas. In the model, the network goes from an unstructured low-activity state to a selective high-activity state after stimulus presentation. This is because the network has different stable attractors before and after stimulus presentation, which is achieved by an untuned attentional input to all neurons (marked as “Attention” in Fig 1E) (Itskov et al., 2011). In addition, we modeled NB stimulation (Fig 1E, right panel) as a non-specific increase in the excitability of the excitatory neurons in the circuit through slight up-regulation of feedforward inputs to the network.

Figure 2 depicts the relationship between the feedforward input and the network’s tuning in the bump state for two neuronal input/output (I/O) functions. When the neuronal I/O function is non-saturating (Fig 2A), the network’s tuning increases monotonically with the input (Fig 2B). Figure 2C plots the population tuning curves for two values of the feedforward input. As the input increases, neurons at the center of the bump increase activity, inhibiting neurons on the edge and sharpening tuning. This appears qualitatively inconsistent with the observation that NB stimulation (modeled here by an increase in feedforward input) results in broader PFC tuning (Qi et al., 2021). However, tuning becomes nonmonotonic in the feedforward input when the neuronal I/O function saturates (Fig 2D). It decreases with the external drive for sufficiently large inputs (Fig 2E). Figure 2F plots the network’s population tuning curves for two simulated conditions: NB OFF and NB ON. NB ON differs from NB OFF by an increase in the strength of the feedforward input that simulates the cholinergic activation. As the neurons at the center of the bump reach saturation, neurons on the edges increase their activity, and the tuning broadens. Therefore, the saturated continuous bump attractor model can display population responses qualitatively similar to the neural responses observed in PFC following NB stimulation (Qi et al., 2021).

To better understand the network mechanisms underlying the balance between enhanced cognitive function and reduced tuning under neuromodulation, we analyzed the behavior generated by our model in repeated network simulations. Specifically, we investigated how changes in network excitability, modeled through changes in the strength of the feedforward input, affected network performance. In the bump attractor model, errors can arise from the diffusion of the bump’s center. This can occur due to stochastic fluctuations in the net inputs into the neurons that change randomly and might shift the bump’s center of mass. We assessed the relationship between feedforward input strength and the diffusion of the bump’s location from 1000 simulations with different stochastic noise realizations (Fig 3A). In the bump attractor model with saturation, we found that the diffusivity of the bump decreases in an intermediate range of feedforward inputs (Fig 3B). Figure 3C shows the distributions of bump centers around the presented targets at the end of the simulated trials. The bump center distribution narrows for larger feedforward inputs, indicating more precise storage of the target’s location and better network performance. These findings are consistent with the general improvement in behavioral performance following NB stimulation (Qi et al., 2021). This shows how, in a bump attractor model of this spatial WM task, reduced neural tuning (Fig 2F) can be associated with improved performance (Fig 3C), provided neurons have saturating responses. Increased performance is explained by a reduction of bump diffusion upon network depolarization. For low feedforward inputs, neuronal responses are not saturating, and the increase in network performance occurs parallel to neural tuning enhancement (Fig. 3B).

This insight from the computational model led us to examine whether the improvement in the behavioral performance of the monkeys after NB stimulation (Qi et al., 2021) could be attributed to a reduction in diffusive errors, i.e., an increase in memory precision. To test this hypothesis, we analyzed saccadic endpoint distributions in control and NB-stimulated trials in the behavioral data of (Qi et al., 2021). In the dataset, responses exhibit two distinct sources of errors (Fig 3D). The first, the “response bias,” is a characteristic offset between the mean response to a fixed target stimulus and this target, and it quantifies the memory accuracy. The second, the “response variance,” measures memory precision computed as the dispersion of the reported locations around the mean response. The continuous bump attractor model predicted that this response variance should be reduced following NB stimulation. Figure 3E-F shows the distribution of response variance in control and NB-stimulated trials, i.e., saccadic endpoints corrected from response biases. Remarkably, we found that the distribution of these corrected reported locations was significantly sharper in NB-stimulated than in control trials for both Remember 1st and Remember 2nd trials (corrected saccades: standard deviations 6. 09 ± _0_. 27° and 5. 0 ± 0. 23° for control and NB stimulation in Remember 1st, Levene’s test *F* = 44. 75, *p* = 2. 68 10^−11^; 7. 31 ± _0_. 35° and 5. 49 ± _0_. 28° for Remember 2nd, Levene’s test *F* = 68. 18, *p* = 2. 38 10 ^−16^, errors are 95% confidence intervals; regression analysis of squared corrected saccade: β _*NB*_ =− 12. 2 *deg*^2^, *p* = 0. 0005,β _*NB/Task*_ =− 12. 4 *deg*^2^, *p* = 0. 014, see Materials and Methods). This suggests that a reduction in diffusive errors could explain the observed improvement in task performance. Our model explains the counterintuitive performance enhancement seen in the behavioral data: reduced neuronal selectivity can still be associated with improved performance if a reduced bump diffusion characterizes network dynamics during the delay period. Our simulations show this is a reasonable condition in bump attractor network models (Fig 2F and 3C).

Next, we investigated if our model could explain how performance modulation by NB stimulation depended on how far S2 was presented from S1 (Fig. 1B; Qi et al., 2021). In our network simulations, we tested the effect of distractor proximity on bump dynamics (Fig 4A-C). Distractors have been reported to attract memory bumps toward their locations (Herwig et al., 2010; Almeida et al., 2015; Lorenc et al., 2018). We found a similar attractive effect in the simulations. Still, these distractor biases did not differ in NB OFF and NB OFF trials (Fig 4B). Moreover, we found that distraction also affected bump diffusion. In particular, nearby distractors (± 45°) increased response variance compared to far distractors (± 90, ± 18_0_ degrees). This was particularly accentuated in NB ON trials (Fig 4C). This pattern was also reflected in the saccadic responses of monkeys. Firstly, distractors biased the distribution of saccadic errors in control and NB-stimulated trials similarly (Fig 4E, regression analysis, β_*NB*_ =− 0. 54°, p=0.357, see Materials and Methods). Secondly, close distractors (± 45°) increased response variance similarly for NB-stimulated and control trials, with comparable bias in both conditions (Fig 4F, regression analysis, β _*NB*_ =− 2. 66 *deg*^2^,*p* = 0. 723, see Materials and Methods). Finally, response variance was significantly lower in NB-stimulated trials than in control trials for far distractors (±90, ±180 degrees, Fig 4F, regression analysis, β _*NB/distance*_ =− 23. 10 *deg*^2^, *p* = 0. 011, see Materials and Methods). This suggests that a specific reduction in bump diffusion in distant distractor conditions underlies improved task performance following NB stimulation.

### Inhomogeneous attractor models

Having shown that the findings of (Qi et al., 2021) are compatible with a simple continuous attractor network model formulation, we wondered about the generality of our results in more realistic network realizations. There are theoretical arguments to expect possible differences. It has been shown that bump diffusivity in an attractor network depends directly on the noise magnitude and the spatial correlations of the inputs and inversely on bump strength (Kilpatrick and Ermentrout, 2013; Krishnan et al., 2018). In the continuous bump attractor network model with neural saturation, NB stimulation modeled as an increase in cellular excitability does not significantly affect the magnitude of noisy fluctuations because recurrent currents in all-to-all, homogeneous networks contribute little noise compared with external inputs. Thus, the primary effect of depolarization in this network is increased bump size, which promotes inertia against diffusion (Kilpatrick and Ermentrout, 2013; Krishnan et al., 2018). However, increased excitability in more realistic network implementations where neurons are connected inhomogeneously to their neighbors may result in changes in the properties of noisy inputs or in distortions of the attractor landscape (Tsodyks and Sejnowski, 1995; Zhang, 1996; Seung et al., 2000; Renart et al., 2003; Itskov et al., 2011; Kilpatrick and Ermentrout, 2013; Kilpatrick et al., 2013), and induce qualitatively different memory diffusion properties.

Indeed, Figure 5 shows how structural heterogeneities impact the diffusion of the bump. Diffusivity decreases markedly with heterogeneity in the network connectivity (Fig 5A) (Kilpatrick and Ermentrout, 2013; Kilpatrick et al., 2013). Figure 5B left (resp. right) plots the time course of the bump’s center for different network initializations of the homogeneous (resp. heterogeneous) bump attractor network. Strong heterogeneities modify the diffusion properties of the bump. This is because, in the presence of heterogeneity, the network undergoes symmetry breaking: the attractor goes from continuous to discrete with only a few stable fixed-point locations (Fig 5B right) (Zhang, 1996). At the fixed point, the bump diffuses weakly in a small basin of attraction instead of being free to diffuse on the entire ring attractor. Nevertheless, even in the presence of strong heterogeneity, tuning and diffusivity of the bump are nonmonotonic, with changes with the feedforward input (Fig 5C-E) qualitatively similar to the homogeneous attractor model, albeit much weaker for the case of diffusivity.

The marked impact that connection heterogeneities have on memory diffusion made us consider how more realistic networks might respond to changes in excitability. In these networks, the conditions to observe lower memory diffusion following increased excitability may depend on the specific biophysical mechanisms that help maintain a quasi-continuous attractor dynamical regime (Stein et al., 2021). We investigated these mechanisms in a biologically realistic model of spiking neurons. The model consists of two populations - excitatory (E) and inhibitory (I) - of strongly recurrent leaky integrate-and-fire neurons with tuned connections (Van Vreeswijk and Sompolinsky, 2005). We assumed that the excitatory-to-excitatory synapses facilitate (Markram et al., 1998; Tsodyks et al., 1998). Facilitation has been shown to promote persistent activity states with highly heterogeneous neural activity (Mongillo et al., 2012; Hansel and Mato, 2013). The network exhibits multistability with an unstructured baseline and structured persistent states (Fig 6B). Additionally, short-term synaptic facilitation has been shown to significantly improve the robustness of WM by slowing down the drift of bump activity in heterogeneous networks (Itskov et al., 2011; Hansel and Mato, 2013). In our current context, we reasoned that short-term plasticity might serve as a potential synaptic saturation mechanism to regulate network activity and tuning in responses to changes in excitability, such as those induced by NB stimulation.

**Figure 6:**
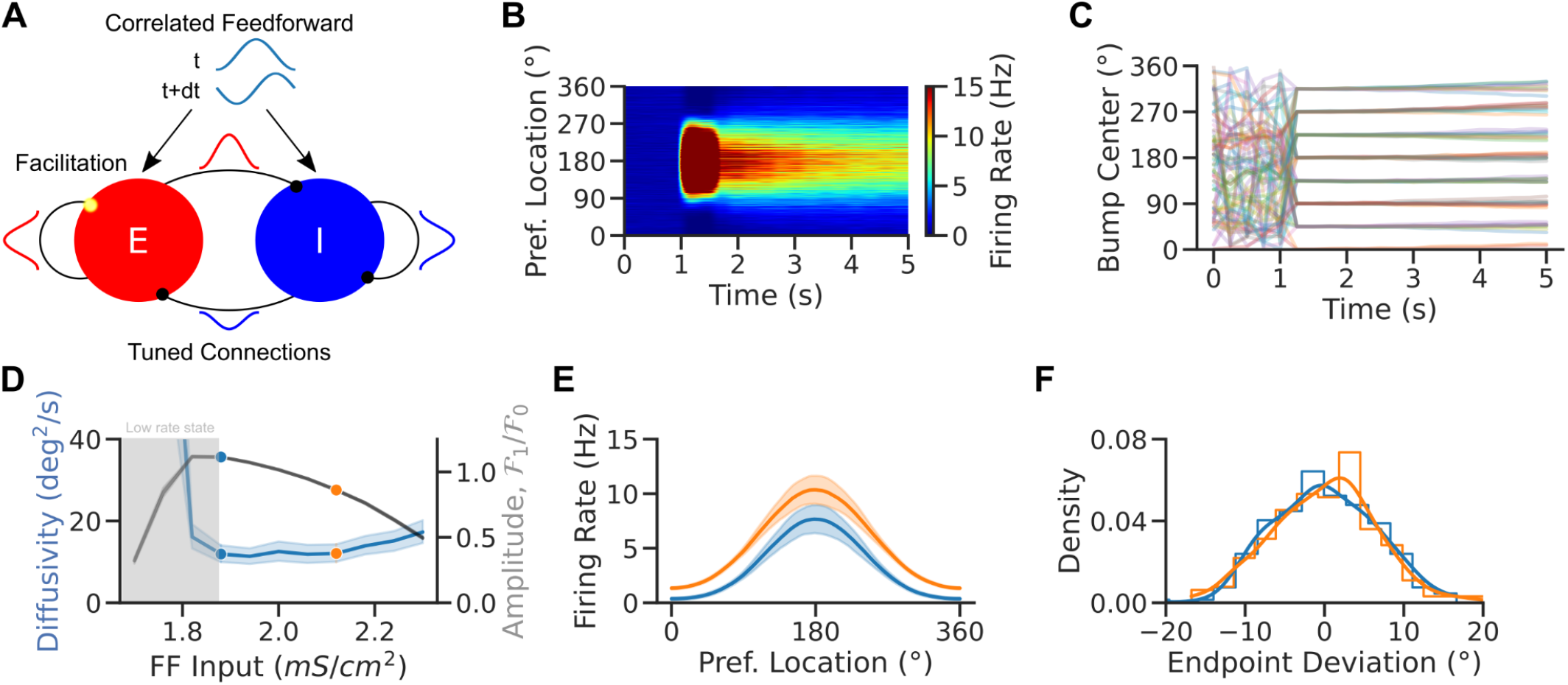
Enhanced excitability weakens network tuning but weakly impacts bump diffusion in spiking attractor networks. A. Network scheme. Spiking EI network with tuned connections (ring model) and facilitating E-to-E synapses. Neurons receive a weakly tuned FF input with a random phase at each time step to induce spatial correlations. B. Local average firing rate vs. time vs. preferred location for one trial in the NB OFF condition. C. Population activity center of mass in 30 simulations for 8 different cue locations. D. Diffusivity (blue) and tuning amplitude (black) vs. FF Input at time t=5 s in the simulations of panel C (averaged over 8 cues and 60 random initializations per cue). E. Population tuning curves averaged over trials at the end of the delay period (tuning curves were recentered before averaging). Increased excitability leads to weaker tuning in the NB ON condition. F. Endpoint deviations (mean corrected endpoint). Colors match dots in D. Lines are kernel density estimates. Error bands 95% confidence intervals.

**Figure 7:**
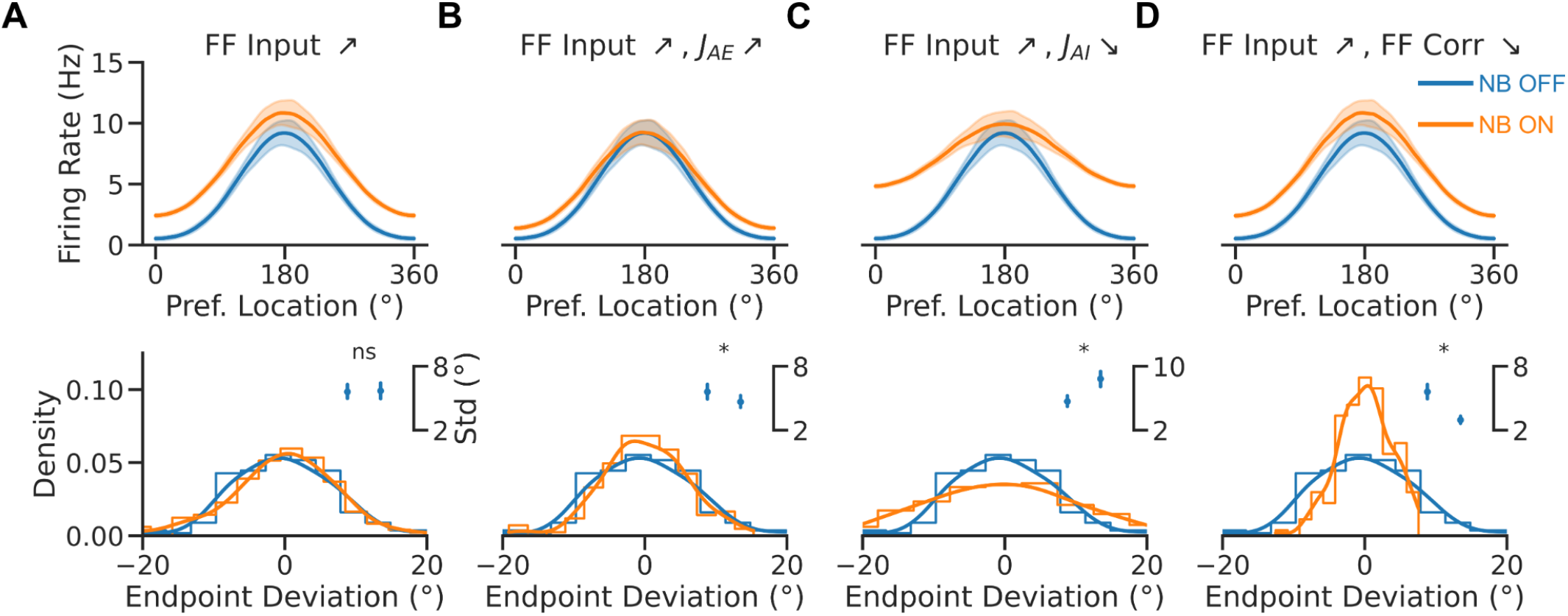
Cholinergic synaptic neuromodulation impacts bump tuning and diffusion, while cholinergic input decorrelation decreases bump diffusion. A-D Top, Population tuning curves (averaged over 8 cues after recentering, with 20 trials per cue) at the end of the delay period. Bottom, Endpoint deviation (mean corrected endpoint). Blue: NB OFF condition (*J* _*E*0_ = 1. 95 *mS cm*^−2^). Orange: NB ON condition (*J* _*E*0_ = 2. 24 *mS cm*^−2^). The NB ON condition has additional modulations in B-D: in B *J* _*EE*_ and *J* _*IE*_ are doubled; in C *J* _*EI*_ and *J* _*II*_ are decreased by 20%; in D FF correlation modulation, *C* _*E*0_, is divided by half. Inset, response Std in the two conditions. Error bars are 95% bootstrapped confidence intervals.

In a strongly recurrent framework, the network dynamically evolves into a state where strong excitation is balanced by strong inhibition such that the *net* input into the neurons is comparable to their thresholds (van Vreeswijk and Sompolinsky, 1996, 1998). Neurons exhibit temporally irregular spiking patterns and heterogeneous firing rates and tuning. However, diffusion in the bump state is particularly weak, as previously observed in sparsely coupled spiking networks (Hansel and Mato, 2013; Stein et al., 2021) and consistent with the effect of heterogeneity in rate-model networks (Fig 5; Kilpatrick et al. 2013; Kilpatrick and Ermentrout 2013). To recover a sizable amount of diffusion, we assumed that the network receives spatially correlated feedforward inputs that simulate inputs from other areas (Fig 6A). As a result of these correlated inputs, the network exhibited weak spatiotemporal activity patterns during baseline activity, and self-sustained bumps in the network showed pronounced memory diffusion, consistent with experimental observations (Wimmer et al., 2014; Stein et al., 2020).

We performed extensive simulations to investigate how parameters affect the model’s dynamics. Figure 6B shows population activity through a single simulated trial for parameters for which the network is multistable: The network goes from an unstructured low rate baseline state to a structured high rate bump state after stimulus presentation. In the bump state, bumps exhibit little drift and diffuse around the cue’s location (Fig 6C). For short simulations (5s in our case), the network exhibits transient tuning in the low rate state (Fig 6C). We investigated how network excitability affected the bump tuning and the bump diffusion in this network by systematically varying the strength of feedforward tonic inputs onto excitatory neurons (Fig 6D). Bump tuning and diffusivity changed with excitability in a qualitatively similar way as observed in the simpler continuous attractor rate model (Fig 3B). Short-term plasticity is the saturation mechanism leading the spiking model to nonmonotonic bump width and diffusion in a range of FF inputs. However, diffusivity varied only weakly in the bump state of the network, even when tuning changed significantly. As a result, also for this more biophysically detailed network model we observed a broadening of population tuning curves upon increasing network excitability (Fig 6D), but this was not accompanied by a significant difference in the distributions of bump center endpoint locations corrected for response biases (Fig 6E). This is consistent with the sharp reduction in endogenously generated bump diffusion caused by network heterogeneity (Fig 5). In realistic EI networks, the mere depolarization of the circuit is unable to reproduce the increased memory precision that would explain task performance improvements upon NB stimulation, which may thus depend on additional cholinergic effects in the circuit.

Modeling cholinergic neuromodulation solely as an increase in excitatory neuron excitability neglects important impacts of acetylcholine release in cortex. Indeed, acetylcholine has been reported to intricately affect multiple cortical mechanisms (Picciotto et al., 2012; Thiele et al., 2012; Minces et al., 2017; Colangelo et al., 2019). We selected a set of specific mechanisms with particular prevalence in the neocortex to test our network model. In particular, in addition to the cellular depolarization of pyramidal neurons that we modeled so far (McCormick and Prince, 1986; Haj-Dahmane and Andrade, 1996), acetylcholine modulates cortical function through: 1) increased synaptic excitatory transmission (Nuñez et al., 2012; Fernández de Sevilla et al., 2021); 2) Reduced GABA release (Salgado et al., 2007; Nuñez et al., 2012); and 3) reduced neuronal correlations (Thiele et al., 2012; Minces et al., 2017).

Therefore, we investigated how, in addition to a change in excitability, changes in synaptic strength or correlations impacted tuning and diffusion in the spiking network model (Fig 7). In the model, increasing excitatory synaptic strength (to both excitatory and inhibitory neurons) did not affect bump tuning but instead significantly reduced bump diffusion (Fig 7B, inset). The reduction of inhibitory synaptic strengths had instead the opposite effect, enhancing bump diffusion (Fig 7C, inset). Finally, reducing spatial correlations in the FF input in the NB ON condition led to comparable tuning and a marked reduction in bump diffusion (Fig 7D, inset). We conclude that the interplay between strengthened excitatory synaptic transmission, diminished neuronal correlations, and increased neuronal excitability, reflect the cholinergic modulations responsible for Qi et al.’s neural and behavioral observations following NB stimulation (Figure 3E-F).

Our simulations, thus, support the association between reduced neuronal tuning and improved memory precision based on attractor dynamics, which is the basis of our interpretations in the bump attractor model simulations. In addition, the model proposes a role for the cholinergic reduction of neuronal correlations in achieving behavioral improvements in WM.

In summary, bump attractor dynamics in inhomogeneous ring networks with short-term plasticity, which effectively causes saturation in recurrent inputs, naturally explain the contrasting effects of NB stimulation on monkey behavioral and neural responses: response variance is generally reduced, while neural tuning broadens.

## Discussion

Here, we show how bump attractor models can reconcile the dynamics of PFC neurons and the behavior of monkeys performing a visuospatial WM task under cholinergic neuromodulation (Qi et al., 2021). We explain how behavior correlates with PFC neural data: cholinergic neuromodulation of PFC reduces neural tuning while reducing bump diffusion, thus enhancing decoding precision and behavioral performance in the task. We found that network depolarization can lead to broader tuning and reduced diffusion in bump attractor models with cellular or synaptic saturation, which can be enhanced by other synaptic and population effects associated with acetylcholine. Our results provide a mechanistic understanding of how elevated prefrontal excitability, possibly combined with other cholinergic mechanisms, impacts spatial WM through changes in bump attractor dynamics in PFC. Our model interpretation is strengthened by the match with the specific distractor condition dependencies observed by (Qi et al., 2021). NB stimulation improved or impaired WM depending on the distance between the target and the distractor. Our model associates impairments in nearby distractor filtering with a broadening of the bump activity that is not counteracted by reduced bump diffusion upon network depolarization. Conversely, increased performance for distant distractors results from robust reductions in bump attractor diffusion.

### Nonmonotonic cholinergic neuromodulation of prefrontal neurons

The effects of acetylcholine on neuronal activity have been investigated in non-human primates with micro-iontophoresis and systemic drug administration. Cholinergic agonists generally increase the activity of prefrontal neurons (Yang et al., 2013; Sun et al., 2017; Dasilva et al., 2019). Conversely, systemic administration of the muscarinic antagonist scopolamine reduces prefrontal activity (Zhou et al., 2011), as does micro-iontophoresis of muscarinic and nicotinic-α7 inhibitors (Yang et al., 2013; Major et al., 2015; Galvin et al., 2020b). However, the increase in activity by agonists is selective for the preferred location of the neuron so that tuning is enhanced, in contrast to the effect of NB stimulation observed by (Qi et al., 2021). This discrepancy may be resolved by noting that the sharpening of prefrontal neurons’ tuning takes place with low doses of cholinergic agonists, while high doses of carbachol or M1R allosteric inhibitors lead to broader tuning in WM tasks (Major et al., 2018; Vijayraghavan et al., 2018; Galvin et al., 2020b). The continuous bump attractor model provides a mechanistic explanation for this nonmonotonic relationship between neural tuning and cholinergic modulation. Figure 2E shows that progressive depolarization of the network initially renders neural tuning sharper but then makes it progressively broader when neurons start engaging the saturating part of their input-output function. Under this framework, intrinsic cholinergic neuromodulation through the stimulation of NB represents a strong release of acetylcholine that mimics the effect of high-dose agonists. A direct empirical exploration of the nonmonotonic dependence predicted by our model could be achieved with graded activation of NB, possibly using optogenetic approaches.

### Cholinergic neuromodulation of prefrontal circuits and performance

The decrease in selectivity by cholinergic overstimulation (Major et al., 2018; Vijayraghavan et al., 2018; Galvin et al., 2020b) has been assumed to correspond to the descending section of an inverted-U function, representing a regime over which cholinergic agonists would impair performance (Galvin et al., 2020a). This interpretation is based on robust behavioral results with drugs targeting the dopamine system, showing parallel inverted-U dose-response of dopaminergic drugs at the behavioral and neural levels (Arnsten, 2011). However, the behavioral impact of cholinergic drugs needs to be clarified. Available evidence from the manipulation of intrinsic acetylcholine release suggests that WM performance may depend monotonically on acetylcholine prefrontal concentration: prefrontal cholinergic input depletion leads to selective WM impairment (Croxson et al., 2011), while electrical stimulation of NB leads to WM performance enhancement (Blake et al., 2017; Liu et al., 2017; Qi et al., 2021). Our network model is consistent with this apparent monotonic relationship. Still, it predicts that a finer sampling of cholinergic modulation of prefrontal circuits should reveal a nonmonotonic component in the behavioral dose-response curve (Fig 3B). Notably, our network simulations in Figure 3B show that the nonmonotonic dose-response relationships for neural tuning and behavior are not aligned in their maximum/minimum, thus breaking the intuition that neural tuning maps directly to behavioral performance.

### Relationship between neural tuning and behavioral performance

Sharper tuning is typically associated with increased performance. This is particularly clear for conditions in which decoding precise stimulus information is essential, as in discrimination tasks after perceptual learning (Li et al., 2004; Yang and Maunsell, 2004a; Raiguel et al., 2006a; Sanayei et al., 2018a). Theoretical studies have shown, however, that broader tuning curves can produce either worse or better performance depending on the task and noise conditions in the network (Pouget et al., 1999; Zhang and Sejnowski, 1999a; Butts and Goldman, 2006; Ma et al., 2006a). For conditions in which stimuli are highly discriminable, and performance depends on the ability to filter distractors or other intervening noise during a memory period, our modeling shows that broader prefrontal tuning can be associated with better memory performance. The connection between reduced neural tuning and increased WM precision depends on specific network implementations and may not generalize to other behavioral readouts in WM tasks. Indeed, previous studies have investigated how improved selectivity can lead to better performance in WM tasks (Yang and Maunsell, 2004b; Compte and Wang, 2005; Raiguel et al., 2006b; Busse et al., 2008; Sanayei et al., 2018b). A network far from saturation shows this tendency. Optimizing mechanisms to increase signal durability during delays may occur at the cost of accuracy, and a stability-accuracy tradeoff may determine performance in a task. Unlike the effects of perceptual learning, learning to perform a WM task also induces a broadening of neural selectivity (Qi et al., 2011; Qi and Constantinidis, 2013).

Based on our model, we propose that NB stimulation leads to reduced bump diffusion in PFC, resulting in a more precise neural code at the end of the delay. This could be tested with large population recordings in PFC during this task. We predict contrasting results of decoders trained on the activity of simultaneously recorded neurons compared to decoders trained on the activity of pseudo-populations of neurons. Decoders trained on pseudo-populations of neurons pooled from different sessions would show reduced accuracy after NB stimulation, likely reflecting the broadening of neural tuning. In contrast, decoding from a true population of simultaneously recorded neurons should show higher accuracy in NB-stimulated trials due to lower levels of correlated noise within the population.

### Circuit mechanisms controlling memory diffusion and neural tuning

Weaker tuning in our continuous attractor network framework can lead to better performance when it is concomitant with a reduction of the noisy diffusion of the memory traces (Pouget et al., 1999; Zhang and Sejnowski, 1999b; Butts and Goldman, 2006; Ma et al., 2006b; Stein et al., 2021). How these two components are integrated into attractor dynamics depends on specific network implementations. We explored here several different network implementations, which have been extensively studied in the literature in relation to the formation of bump attractors in rate models (Amari 1977, Wilson Cowan 1972) and in spiking models (Compte et al 2000; Hansel and Mato 2013), diffusion of the bump (Zhang 1996; Kilpatrick and Ermentrout 2013; Compte et al 2000; Wimmer et al 2014; Krishnan et al. 2018), sensitivity to distractors (Compte et al 2000), or the impact of heterogeneities (Hansel and Mato, 2013; Kilpatrick and Ermentrout 2013; Kilpatrick et al 2013; Seeholzer et al). We specifically asked how neural excitability jointly affects bump diffusion and tuning in these models. In idealized homogeneous networks, the mere depolarization of the network close to saturation provides both broader tunings and reduced diffusion; in heterogeneous networks, however, additional mechanisms are required to observe a significantly reduced bump diffusion. Our modeling identifies a reduction in spatial correlations as one possible additional mechanism. Indeed, changes in neuronal correlations significantly impact the diffusion of bump attractors (Kilpatrick and Ermentrout, 2013), and bump diffusion is a determinant of behavioral imprecisions in these tasks (Wimmer et al., 2014). A role for modulation of neuronal correlations in our data is plausible since cholinergic activation is known to reduce neuronal correlations in multiple brain areas (Thiele et al., 2012; Minces et al., 2017). Furthermore, a reduction in correlated variability between neurons has been linked to increased attention (Cohen and Maunsell, 2009; Mitchell et al., 2009; Herrero et al., 2013), typically associated with improved cognitive performance.

We do not view our modeling approach as a commitment to a specific mechanistic basis for neuromodulating neuronal correlations. Previous computational work has explored the effect of feedforward excitation on correlations in recurrent E-I networks (Rosenbaum et al., 2017), showing that fluctuations in the feedforward inputs can drive correlated activity. Here, we implemented a similar mechanism but did not explore in depth the possible role of recurrent dynamics on the neuromodulation of neural correlations. Indeed, (Darshan et al., 2018) have shown that recurrent structural motifs can generate collective network activity. How specific pathways of neuromodulation might impact correlations induced by recurrent circuitry remains an open question.

### A methodological approach to identifying mechanisms of behavioral alterations

The challenge of relating cellular or synaptic mechanism modulations with changes in cognitive function using biological neural network models has been pointed out before (Stein et al., 2021). To constrain network models, obtaining neural-level information about how the considered cellular changes affect neural circuits implicated in the behavioral readout is crucial. We applied this approach by constraining our models with neural tuning curves obtained with and without NB stimulation and testing their behavioral predictions. This has allowed us to identify a role for neural saturation dynamics, excitatory synaptic transmission, and neural co-variability in determining cholinergic WM improvements. This approach is necessary to advance toward a circuit-level understanding of how alterations in neuromodulators and receptors underlie cognitive dysfunctions, so efforts must be put into generating relevant neural-level data upon hypothesized cellular and synaptic perturbations.

In sum, we show that the bump attractor model can provide a causal link between PFC electrophysiology and the complex pattern of behavioral improvement and impairment in WM caused by endogenous acetylcholine release. The relevant mechanisms in these network models are cholinergic cellular depolarization, cholinergic excitatory synaptic enhancement, neuronal saturation, and cholinergic reduction of input correlations. Our evidence supports that attractor dynamics in PFC are under the neuromodulatory control of cholinergic centers to improve cognitive performance in WM.

Table 7: Backbone and full-atom RMSD comparison of TCR CDR3 loop design. A signed Wilcoxon paired two-sided rank statistical test between FrameDiPT and the best AlphaFold model is performed at significance level p-value < 0.05. Underline means significantly different from the best AlphaFold model.

## Acknowledgments

We acknowledge support from the Instituto de Salud Carlos III of Spain (Ref: AC20/00071), the Spanish Ministry of Science and Innovation co-funded by the European Regional Development Fund (Refs: RTI2018-094190-B-I00, PID2021-125453OB-I00), and the CERCA Programme/Generalitat de Catalunya to AM, DB, and AC; and support from the US National Institutes of Health under grant RF1 AG060754 and from the US National Science Foundation under grant CRCNS-2011514 to CC.

The authors declare no competing financial interests

